# A matter of life and death – Hunting tactics of Southern elephant seals *(Mirounga leonina)* and anti-predatory behaviours of their prey

**DOI:** 10.1101/2023.10.19.563063

**Authors:** Mathilde Chevallay, Christophe Guinet, Pauline Goulet, Tiphaine Jeanniard du Dot

## Abstract

For air-breathing marine predators that must save energy during dives, the ability to adopt hunting tactics that minimise the risk of triggering an escape reaction from the prey is crucial for an efficient foraging. Female Southern elephant seals *(Mirounga leonina,* SES hereafter) forage on small mesopelagic prey that they must hunt almost continuously to meet their high-energy requirements. Here we aimed at understanding how these large time-limited deep-divers can efficiently exploit their small sized prey. To do so, we used data recorded by a new biologger, the sonar tag, deployed on SES during their post-breeding foraging trips. This tag combines an active acoustic sensor with ultra-high-resolution movement and bioluminescence sensors, offering a unique opportunity to simultaneously describe SES’ hunting tactics to capture their prey and defence mechanisms employed by prey. We analysed more than 5,800 prey capture events in nine SES and show that they adopt a “stalking” hunting behaviour allowing them to get as close as possible to their prey before attacking. The ability of SES to adopt stealth approach tactics, minimising the risk of initiating a flight reaction from their prey, might be a key factor in the success of this far-ranging generalist predator.

## 1. INTRODUCTION

Predator-prey interactions play a key role in ecosystem structure and functioning. While predators shape prey population dynamics (i.e., survival, growth, behaviour, etc.), prey availability and accessibility regulate predator populations (Estes & Duggins 1995, Frederiksen et al. 2006, Letnic et al. 2012). Fine-scale interactions between predators and their prey, i.e., how predators find, select, and capture their prey, and alternatively how prey detect predators, and react to imminent predation, are key factors in determining both predators’ hunting efficiency and prey survival (Cooper Jr 1997, McHenry et al. 2009, Stewart et al. 2013). On one hand, to avoid capture, prey can flee, fight back or stay still depending on hunting tactics of their predators, distance from their predators, and accessibility to a shelter (Broom & Ruxton 2005, Eilam 2005). On the other hand, predators must adopt hunting tactics that increases their chances of successful capture while avoiding excessive energy expenditure. This is particularly true for air-breathing marine predators, which are limited by oxygen availability and must regularly return to the surface to breathe (Thompson 1993). For these predators, active chase must be avoided in order to preserve oxygen reserves and increase time spent diving and searching for prey. Therefore, the ability of air-breathing predators to adopt hunting tactics that limit the risk of triggering an escape reaction from the prey is crucial for an efficient foraging.

Studying fine-scale predator-prey interactions in the marine environment is particularly challenging. Development of animal-borne biologging technologies advanced the study of deep-diving animal foraging behaviour. GPS sensors for example can help identifying the foraging areas of marine animals (e.g. (Hindell et al. 1991, Sommerfeld et al. 2015, Reisinger et al. 2018)), whereas high-resolution head-mounted accelerometers allow the detection of foraging events by measuring rapid head movements potentially associated with prey capture attempts (Suzuki et al. 2009, Viviant et al. 2010, Guinet et al. 2014, Ydesen et al. 2014). Accelerometers also record the movements and postures of animals, providing a fine-scale description of their hunting behaviour (e.g. (Johnson et al. 2008, Shepard et al. 2008, Jouma’a et al. 2016, 2017, Le Bras et al. 2016, 2017). Diet analyses (i.e. stomach contents, scat analyses, stable isotopes analyses, genetic markers) provide an overview of prey consumed during foraging trips (e.g. (Cherel et al. 2008, Jeanniard-du-Dot et al. 2017)) but fine-scale information on prey size and behaviour is still challenging to obtain. Recently developed miniature video cameras have provided the first direct information on prey and predator-prey interactions (Adachi et al. (2021), Naito et al. (2013), Yoshino et al. (2020) on Northern elephant seals *(Mirounga angustirostris)*, Bowen et al. (2002) on harbour seals *(Phoca vitulina)*, Kernaléguen et al. (2016) on Australian fur seals *(Arctocephalus pusillus doriferus)*, Davis et al. (1999) on Weddell seals *(Leptonychotes weddellii)*).

Miniature cameras deployed on Northern elephant seals provided unique and novel information on prey targeted by female elephant seals (Naito et al. 2013, Yoshino et al. 2020, Adachi et al. 2021). As suggested by isotopic analyses (Cherel et al. 2008), video footages confirmed that females mainly target myctophid species, and to a lesser extent mesopelagic squids. Small size of prey relative to the large size of females implies that they must feed continuously to meet their high-energy requirements. This raises the question of how these large time-limited deep-divers can efficiently exploit these small prey. Cameras require a light-source to illuminate dark waters where elephant seals roam, which could potentially alter both predator and prey behaviour. Moreover, high power consumption means these devices can only record selectively, for relatively short periods, during multi-month foraging trips. Therefore, extensi ve deployments are lacking and the way elephant seals can efficiently exploit their tiny prey is still poorly known. In order to have a more complete and detailed picture on predator-prey interactions in deep-diving predators such as elephant seals, we used a miniature sonar tag, combining active acoustic signals with high-resolution movement sensors. This tag uses a high-click rate narrow, forward directed beam that can insonify prey at a distance up to 6 m in front of the animal. Even if data recorded by the tag do not allow for the identification of the species of prey targeted by seals, it provides information on their acoustic size – a proxy for actual size – gregarious and escape behaviours (Goulet et al. 2019). Including high-resolution movement sensors, the tag offer the opportunity to study the hunting tactics of seals and more generally fine-scale predator-prey interactions. A more recent version of the sonar tag also include a high-resolution bioluminescence sensor to study bioluminescent behaviour of prey, especially how they might emit light flashes to defend themselves against predators (Goulet et al. 2020) by temporarily dazzling predators and using the opportunity to escape. Therefore, data recorded by this tag will enable the investigation of questions raised by the results obtained with the camera deployments.

Previous sonar tag deployments on Southern elephant seals demonstrated that seals have surprisingly a strong sensory advantage over their prey, allowing them to detect prey several seconds before striking, well before they react (Chevallay et al. in revision). As Northern elephant seals, they target small mesopelagic prey, which they have to hunt continuously to meet their energy requirements. As they are constrained to return to the surface to breathe, they must be efficient during their time-limited hunting periods. Therefore, we hypothesised that the ability of Southern elephant seals to detect their prey in a distance enabled them to adopt stealth approach tactics to minimise the risk of initiating a flight reaction from their prey, therefore avoiding energy-expensive chase. To test that hypothesis, we took advantage of data recorded by the sonar tag to describe the hunting sequence of elephant seals and to describe the prey defence mechanisms to see how it influences the seals’ capture success.

## 2. MATERIALS AND METHODS

### 2.1. Device deployments and data collection

Data were collected on 9 post-breeding female Southern elephant seals in October 2018 (n = 3), October 2019 (n = 3) and October 2020 (n = 3), in Kerguelen Islands (49°20’S – 70°26’E). Females were captured with a canvas head-bag, anaesthetized with a 1:1 combination of Tiletamine and Zolazepam (Zoletil 100 – 0.7 to 0.8 ml/100 kg) either injected intravenously (McMahon et al. 2000) or using a deported intramuscular injection system. Females were measured to the nearest centimetre and weighted to the nearest kilogram. They were equipped with a neck-mounted Argos tag (SPOT-293 Wildlife Computers, 72x54x24 mm, 119 g in air), a back-mounted CTD tag (SMRU-SRDL, 115x100x40 mm, 680 g in air) and either a head-mounted DTAG-4 sonar tag (n = 7, 95x55x37 mm, 200 g in air, see Goulet et al. (2019) for further details on the device) or a DTAG-4 sonar-light tag (n = 2, 92x69x37 mm, 250 g in air). Tags were glued to the fur using quick-setting epoxy glue (Araldite AW 2101, Ciba). CTD tags recorded conductivity, temperature and depth at 0.5 Hz. Sonar and sonar-light tags were programmed to sample GPS position (up to every minute), tri -axial acceleration (200 Hz), tri-axial magnetometer (50 Hz) and pressure (50 Hz). Sonar-light tags also integrated a light sensor specifically designed to sample bioluminescence events with a sampling frequency of 50 Hz (Goulet et al. 2020). The active sonar within the tags recorded acoustic backscatter returning from 10 µs pings with a centre frequency of 1.5 MHz, at a 25 Hz ping rate for 2018 and 2019 deployments and 12.5 Hz for 2020 deployments. The active sonar operated with a 3.4° aperture beam width and a 6 m range (Goulet et al. 2019). Tags were recovered in late December to January when females returned to shore to moult using the same capture and sedation methods. All experiments were conducted under the ethical regulation approval of the French Ethical Committee for Animal Experimentations (2019070612143224 (#21375)) and the Committee for Polar Environment.

### 2.2. Data analyses

Data recovered from tags were analysed using custom-written codes and functions from www.animaltags.org in MATLAB version 2022b (The MathWorks 2022). Statistical analyses were conducted in R software version 3.5.1 (R Core Team 2018).

#### Prey capture attempts identification

Prey capture attempts (PrCAs hereafter) were detected from the 200 Hz tri-axial acceleration data recorded by the sonar tags, by computing the norm of the differential of the tri-axial acceleration (norm-jerk hereafter), as described in Goulet et al. (submitted). Spikes in the norm-jerk higher than 400 m.s^-2^ were classified as PrCAs. As prey may be encountered in patches or may elude capture, leading to a bout structure in prey strikes, strikes occurring less than 15 s from the previous strike were grouped in the same bout as described by Goulet et al. (submitted).

#### Sonar data analysis

Sonar data recorded during bouts were displayed as echograms, showing the time on the horizontal axis and the distance from the sonar tag on the vertical axis, extending from 10 s before the bout start time to 2 s after the bout end. Because of the large number of bouts detected (ranging from 4595 to 12039 per animal), only a random subsample of 5 to 10% of bouts performed during acoustic data recordings were analysed. Different variables describing predator-prey interactions were extracted manually from echograms following the method described by Goulet et al. (submitted): (1) number of prey, (2) prey evasive behaviour, (3) prey acoustic size, (4) capture success. (1) The number of prey was defined as the maximum number of independent echo traces within a same ping (Jones et al. 2008). It was scored as one, two or more than two, i.e. a school of prey. (2) Prey evasive behaviour was identified from the closing speed between predator and prey, which will vary in case of prey reaction, resulting in a change in the slope of the prey echo trace (Goulet et al. 2019, Vance et al. 2021). If a prey reaction was observed during the bout, prey was considered as evasive. (3) Prey acoustic size was estimated from the −20 dB echo pulse width measured on the widest part of the prey trace on evasive prey only (Burwen et al. 2003). Finally, (4) when prey trace was seen within 50 cm of the sonar transducer within the last second of the bout, capture was considered successful. If prey trace was seen further than 1 m away from the sonar transducer during the last second of the bout or was evident after the bout ended, bout was classified as unsuccessful. All other cases were classified as “unknown” (Goulet et al. submitted).

#### Bioluminescence data analysis

Bioluminescent flashes were detected as spikes in the 50 Hz light data recorded by the sonar-light tags decimated at a 5 Hz sampling rate. Spikes in the data higher than 0.07 (arbitrary unit) were classified as flashes (Goulet et al. 2020). Events detected within less than 5 s of each other were considered coming from the same bioluminescence event (Goulet et al. 2020). If a bioluminescence event started between 5 s before and 5 s after the bout, it was considered associated with the bout. Flash events may consist of a series of short flashes, so the 5 Hz data may not have sufficient resolution to accurately describe them. Therefore, we used 50 Hz light data recorded during flash events to calculate the number of peaks in the light signal as well as the intensity and duration of each peak.

#### Predator approach behaviour

Metrics describing seals’ fine-scale behaviour during the approach and capture phase (i.e. between the 10 s preceding each first strike of a bout (Chevallay et al. in revision) and the last strike of the bout) were extracted from the 200 Hz tri-axial accelerometer data decimated at 5 Hz. Posture of seals was inferred from Euler angles (i.e. pitch angle (rotation around left-right axis), roll angle (rotation around longitudinal axis) and heading angle (rotation around dorso-ventral axis)) (see Johnson and Tyack (2003) for details of the formulas). Bouts were characterised in term of duration (time elapsed during the first and last head strike of the bout) and intensity, defined as the RMS of the norm-jerk signal during the bout. To know whether seals actively swam or passively glided when hunting, we determined the number of flipper strokes between the 10 s preceding each bout and the last strike of the bout. Flipper strokes were detected from dynamic acceleration of the lateral axis (Sato et al. 2007) by applying a high-pass filter with a cut-off frequency of 0.6 Hz, i.e. 70% of the dominant stroke frequency, on the lateral acceleration (Chevallay et al. in revision). Finally, the swimming effort, a proxy of energy expenditure during swimming, was defined as the summed absolute values of the filtered lateral acceleration (Maresh et al. 2004, Aoki et al. 2012). Seals are usually negatively buoyant in the first week of the post-breeding foraging trips (Biuw et al. 2003) and therefore in the period sampled by the sonar tags. Seals are thus expected to glide when descending and swim actively when ascending. To avoid conflating these two distinct movement styles when analysing hunting tactics, bouts were grouped according to the vertical movement prior to each bout. Bouts with an initial depth-rate higher than 0.25 m/s and lower than −0.25 m/s were grouped as, respectively, descending and ascending bouts (Chevallay et al. in revision).

An exploratory analysis of seals’ hunting tactics was first conducted on a sub-sample of bouts by making a 3D animation of seals’ behaviour during the approach phase. This exploratory analysis identified three main approach modes, which differed in bout duration, swimming effort, roll and heading ranges. Therefore, a K-means algorithm with 3 clusters was applied on these behavioural metrics for ascending and descending bouts separately. Each resulting cluster was described in terms of bout duration, bout intensity, swimming behaviour and posture.

### 2.3. Statistical analyses

Approach behaviour between prey types were compared using generalized linear mixed models (GLMM hereafter, R package ‘MASS’, (Ripley et al. 2013)) with gamma distribution, with bout duration (s), bout intensity (m.s^-2^), number of flipper strokes, swimming effort (m.s^-2^), pitch, roll or heading angle extent (°) as response variables, and prey type (schooling vs. single prey; single evasive prey vs. single non-evasive prey) as fixed effect. Individual seal identities were used as random effects.

Influence of prey evasive behaviour on the likelihood of adopting an approach mode was tested using mixed effects logistic regression, with approach mode (either high or low-energy approach) as a binary response variable, flight initiation distance (m) as fixed effect, and individual identities as random effects.

The impact of prey reaction on seals’ capture success was modelled using mixed-effect logistic regression models with capture success as a binary response variable (scored as 1 or 0), reaction type (flash or escape attempt) or timing of reaction as fixed effects and individual identities as random effects.

Finally, the effect of prey characteristics on capture difficulty was tested using GLMM with gamma distribution with the bout duration, the number of head strokes and swimming effort (proxies of prey capture difficulty, (Goulet 2020)) as response variables, and acoustic size or flight initiation distance as fixed effects. Individual seal identities were used as random effects.

For all models, assumptions were checked and any deviation from them were corrected.

## 3. RESULTS

Tags recorded data during 29-79 days. Over that period, seals performed 69 ± 7 dives per day (mean ± sd, min 57 dives per day, max 79 dives per day), and 510 ± 53 bouts per day (range 443-610 bouts per day). Seals performed dives at 413 ± 89 m depth (mean ± sd) for 19 ± 3 min, and spent 10 ± 2 min at the bottom of the dive. A diel diving pattern was observed for almost all individuals, with deeper and longer dives during the day (449 ± 187 m, 21 ± 4 min) than at night (329 ± 70 m, 16 ± 1 min, LMM, P < 0.001).

### 3.1. Prey characteristics

Prey characteristics were inferred from sonar data during 5778 bouts for the nine females equipped with sonar tags between 2018 and 2020. Single prey represented 99.2 ± 0.4% of prey targeted by our studied seals. Among single prey, 81.4 ± 8.1% tried to escape (Table 1). For the two females equipped with sonar-light tags, flashing prey represented 24.3 and 29.5% of their targeted prey. Acoustic size of prey ranged from 5.0 to 9.0 cm (1^st^ and 3^rd^ quantiles, Q1-Q3 hereafter) (min 1.6 cm, max 32.8 cm, Figure 1). We applied a mixture model to the distribution of prey acoustic sizes (R package mixtools (Benaglia et al. 2010)), and identified three size groups (Figure 1): small prey measuring 3 to 6 cm (4.3 ± 1.0 cm), medium-sized prey measuring 6 to 11 cm (7.6 ± 2.0 cm), and large prey measuring more than 11 cm and up to 35 cm (11.8 ± 4.1 cm).

**Figure 1:**
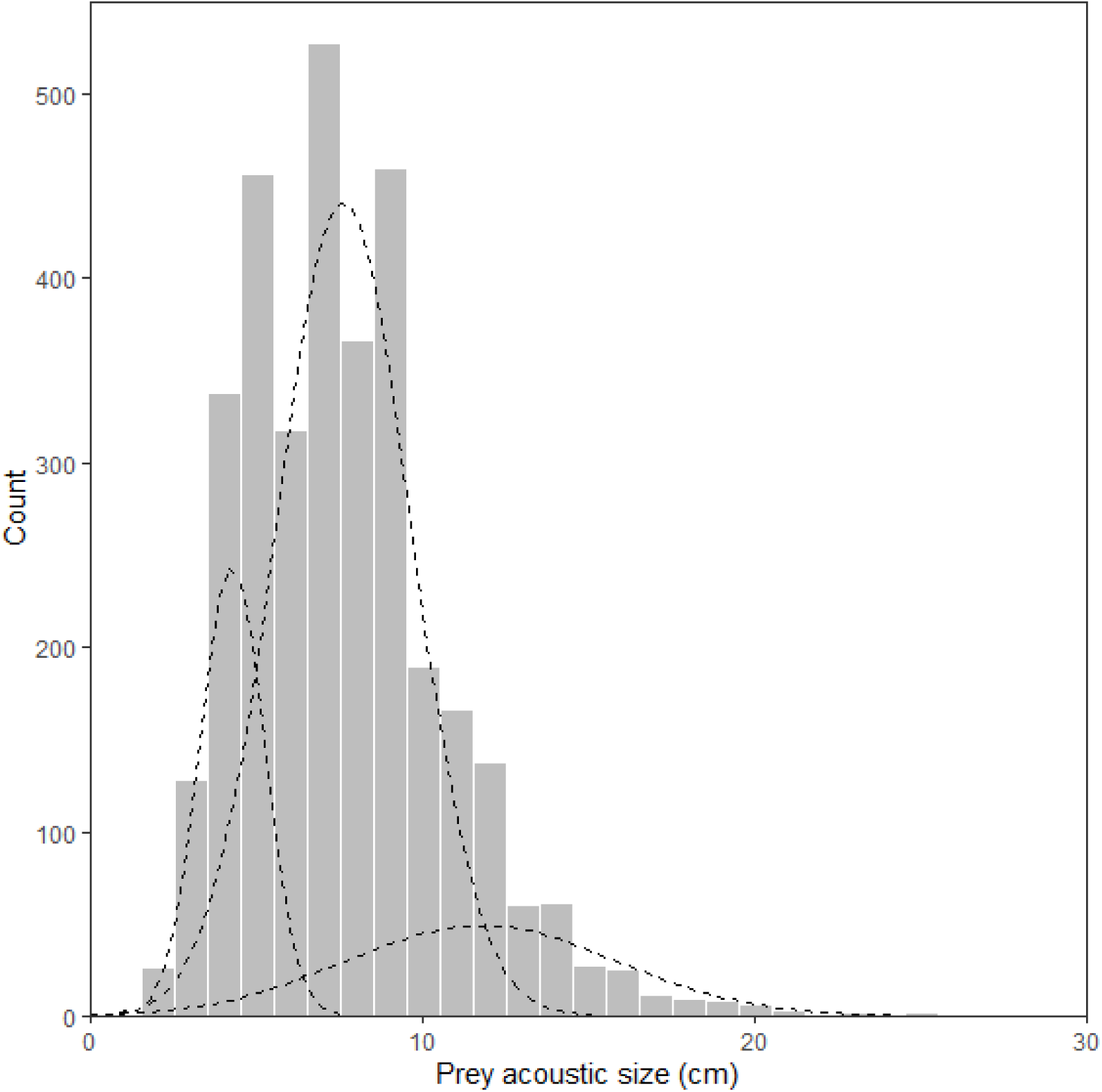
Acoustic size distribution of prey targeted by nine female Southern elephant seals equipped with high-resolution movement and sonar tags in Kerguelen Islands between 2018 and 2020. Prey acoustic sizes were estimated from the −20 dB echo pulse width measured on the widest part of the prey echo trace. A mixture model was applied which identified three normal distributions in the acoustic size data.

**Table 1:**
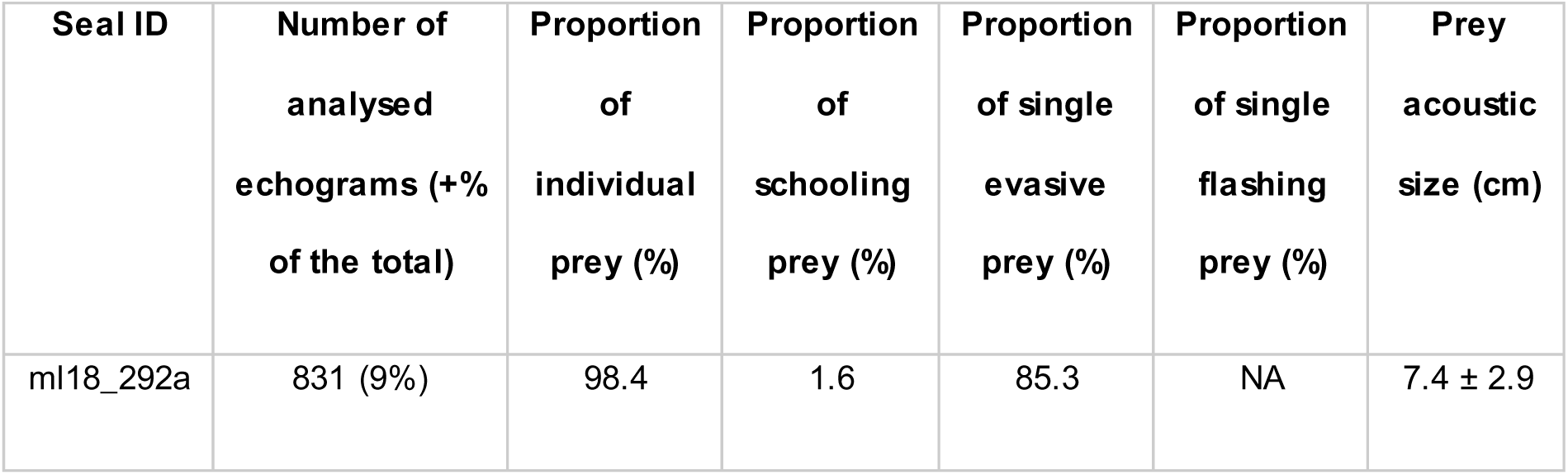
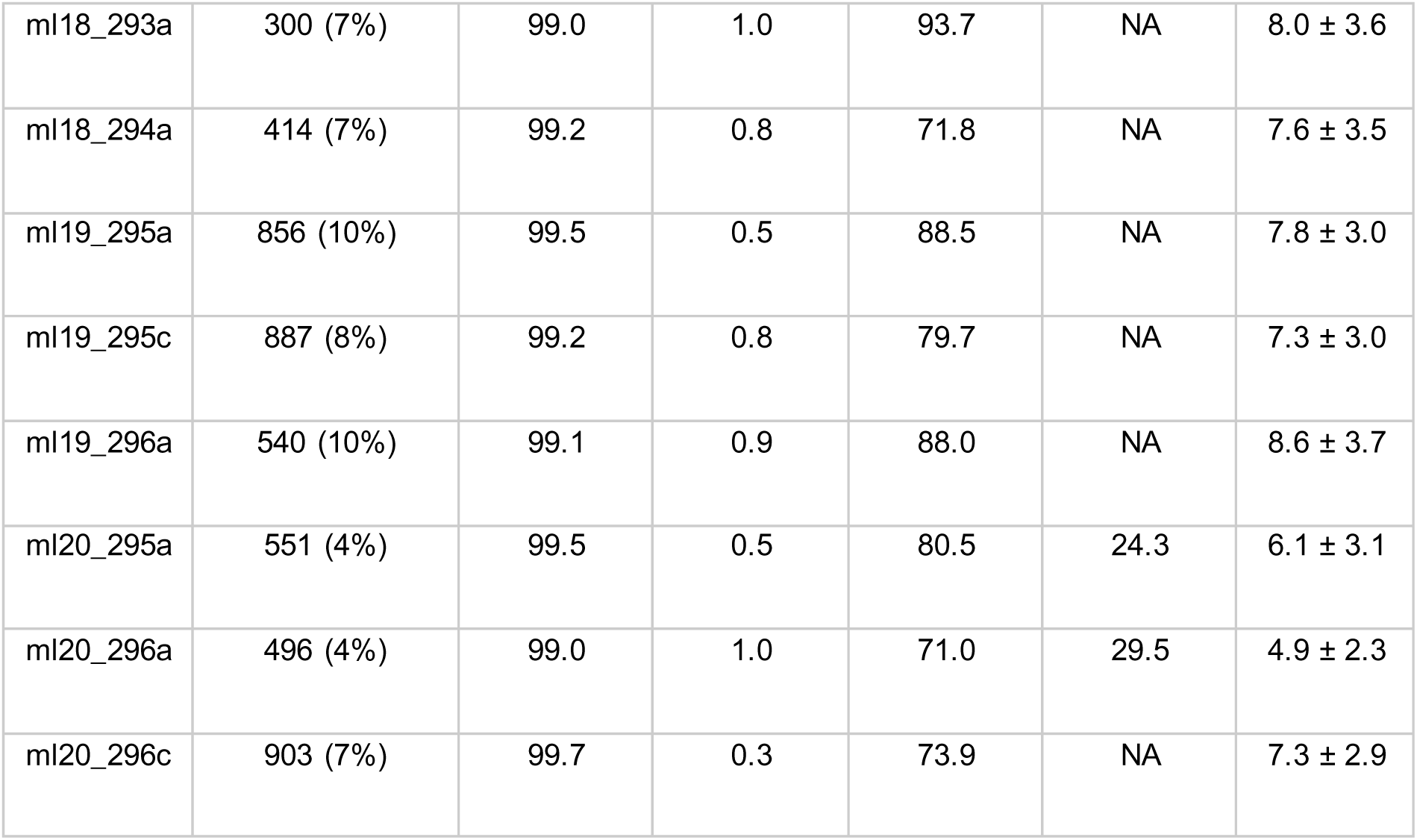
proportion of single prey, schooling prey, evasive prey and flashing prey and prey acoustic size targeted by nine female Southern elephant seals equipped with high-resolution movement and sonar tags in Kerguelen Islands between 2018 and 2020. Seal ID comprises the two first letters of the Latin species binomial followed by the year and Julian day of attachment, and a letter denoting the sequential animal of day.

### 3.2. Predator hunting behaviour

Gliding behaviour preceded 91% and 60% of descending and ascending bouts, with a stop of active stroking 6.7 ± 1.4 s and 5.5 ± 1.6 s before the first strike of the bout (Figure 2). Then, depending on prey behaviour, seals displayed three hunting modes: low-energy mode, associated to non-evasive prey, high-energy mode, associated to evasive prey, and sinuous mode, associated to schooling prey. Low-energy mode was the most frequent in both descents (62% of bouts) and ascents (67% of bouts). Sinuous mode was observed for 27% and 21% of descending and ascending bouts, and high-energy mode was the least frequent mode, observed for only 11% and 12% of descending and ascending bouts.

**Figure 2:**
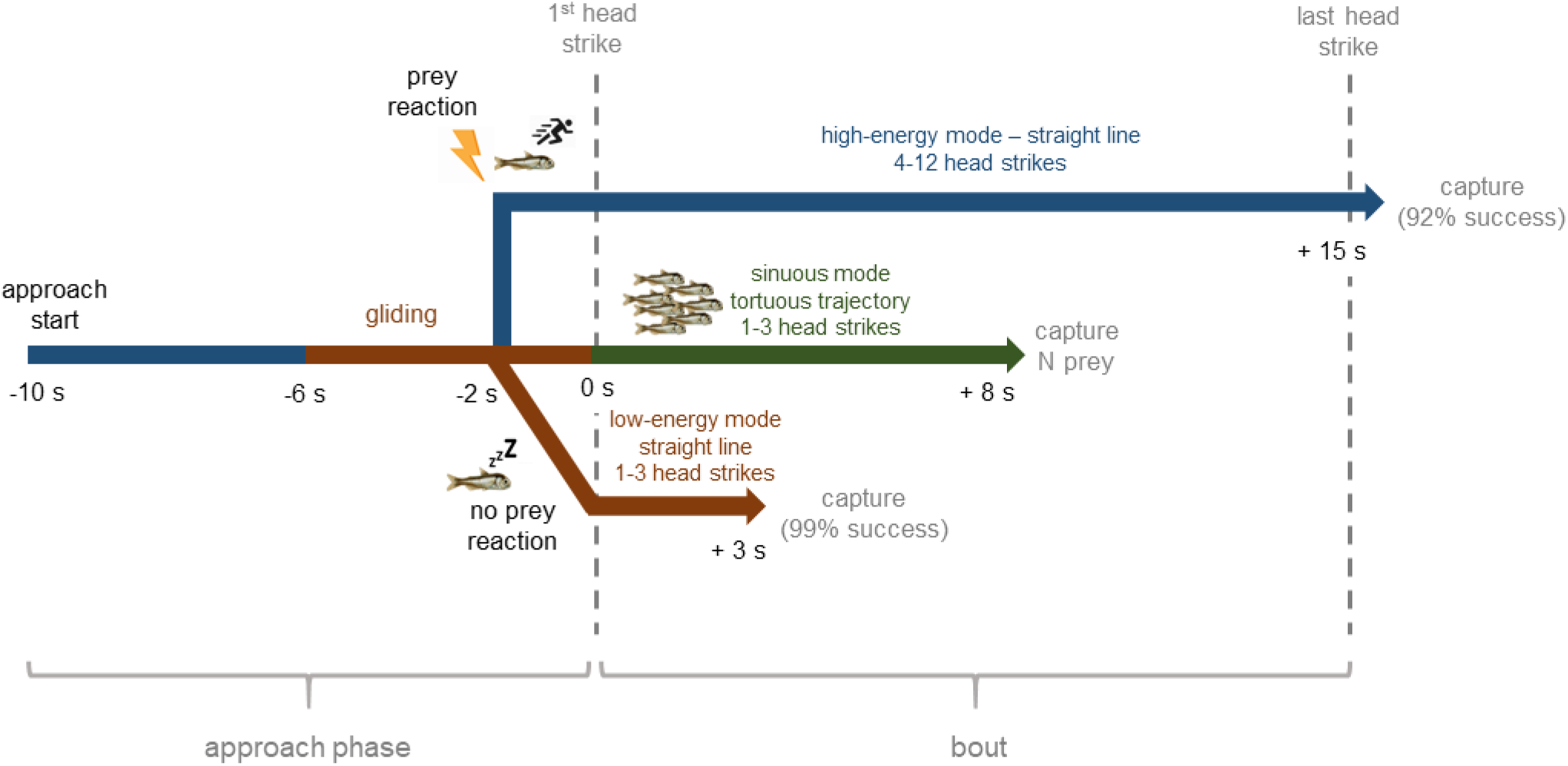
Summary of the hunting sequence of Southern elephant seals. Predator-prey interactions were described on nine females equipped with high-resolution movement and sonar tags in Kerguelen Islands between 2018 and 2020. The hunting sequence was described as the sequence of events from the beginning of the approach phase (i.e. 10 seconds preceding each bout) to the final head strike of the bout. Seal hunting behaviour was derived from accelerometer and magnetometer data, and prey behaviour was inferred from sonar data. Capture success was inferred from sonar data for individual prey only, as we were not able to determine the number of prey ingested in a school.

Low-energy modes were characterised by short bouts of 1 to 3 (Q1-Q3) intense head strikes (406.5 ± 246.5 m.s^-2^, Figure 3) lasting 0.3 to 3.7 s (Q1-Q3), and low swimming efforts (37.2 ± 30.2 m.s^-2^, Figure 3). High-energy modes were characterised by long bouts of 4 to 12 (Q1-Q3) low intensity head strikes (287.7 ± 107.8 m.s^-2^) usually lasting 5.0 to 20.2 s (Q1-Q3) and up to 36.6 s (90^th^ percentile, Figure 2). This mode is associated with the highest values of swimming effort (99.5 ± 83.7 m.s^-2^), especially during ascents (Figure 2). The probability of adopting low or high-energy hunting mode depended on prey evasive behaviour: the greater the flight initiation distance (FID hereafter), the greater the probability of adopting a high-energy hunting mode (mixed-effects logistic regression, P < 0.001, Figure 2). Prey that reacted sooner were associated with longer bouts (LMM, P < 0.001): bout duration (s) = (9.8 ± 2.0) + (8.9 ± 0.7) * FID (m)) with a higher number of head strikes (LMM, P < 0.001, number of head strikes = (6.4 ± 1.0) + (4.8 ± 0.3) * FID (m)) and a higher swimming effort (LMM, P < 0.001, swimming effort (m.s^-2^) = (64.9 ± 10.2) + (41.5 ± 4.1) * FID (m)). Finally, schooling prey were associated with sinuous approaches, characterised by the highest values of roll and heading extent, short bouts of 1 to 3 intense head strikes (435.3 ± 288.4 m.s^-2^) lasting 0.3 to 10.0 s (Q1-Q3), and low swimming efforts (55.2 ± 73.9 m.s^-2^, Figures 2 and 3).

**Figure 3:**
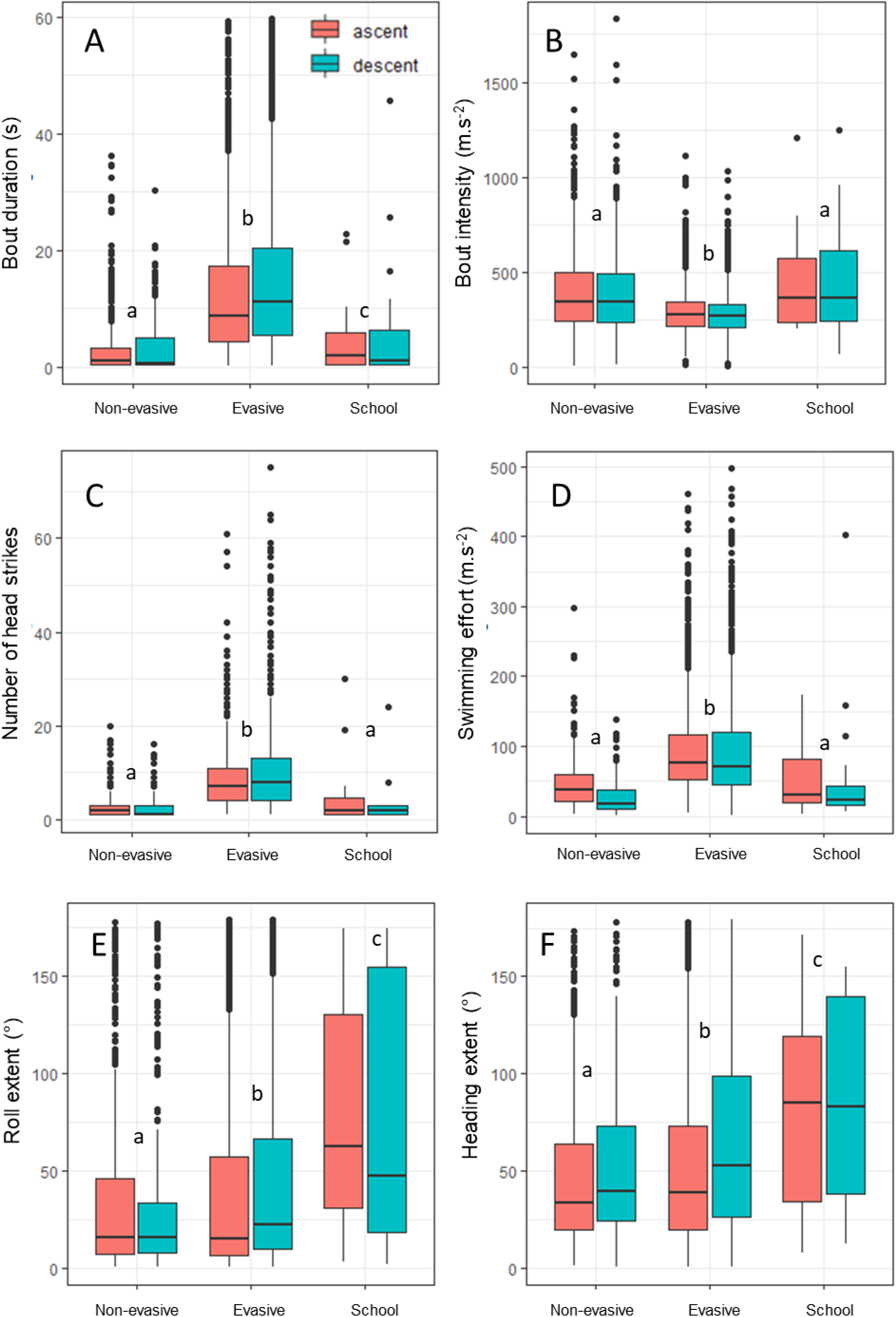
Characteristics of prey-dependant hunting behaviour of nine female Southern elephant seals equipped with high-resolution movement and sonar tags in Kerguelen Islands between 2018 and 2020. Behavioural metrics were derived from accelerometer and magnetometer data recorded between the 10 seconds preceding each bout and the last strike of the bout: ((A) bout duration (s), (B) bout intensity (m.s^-2^), (C) number of head strikes, (D) swimming effort (m.s^-2^), (E) roll extent (°) and (F) heading extent (°)). Letters in superscript indicate significant differences in behavioural parameters between prey types (GLMM, P < 0.05).

### 3.3. Prey anti-predatory behaviours

Anti-predatory behaviours of elephant seals’ prey were studied on 1047 bouts for which data on both evasive and flashing behaviours were available. Escape attempt was the most frequent defence mechanism, detected in 59% of prey, while 19% of prey did not show any reaction, 17% of prey flashed and tried to escape, and 5% only flashed. Prey initiated flight reaction 2.4 ± 2.4 s (Q1-Q3: 3.5 – 0.4 s) before the first strike of the bout, at 0.7 ± 0.4 m (Q1-Q3: 0.5 – 0.9 m) from seals. Prey flashed 1.9 ± 1.5 s (Q1-Q3: 3.3 – 0.5 s) before the first strike of the bout. Flash events usually lasted 1.7 ± 3.0 s (Q1-Q3: 0.2-1.6 s), as a series of 2 ± 2 flashes (Q1-Q3: 1-3 flashes) of 0.17 ± 0.24 s each (Q1-Q3: 0.04-0.19 s) and of 0.18 ± 0.15 intensity (Q1-Q3: 0.10-0.18).

### 3.4. Impact of prey behaviour on seals’ capture success

We estimated prey capture success at 99% when prey did not react to the seals’ approach. Capture success was not significantly different between prey that tried to escape and flashed (95%) and prey that only tried to escape (92%) (mixed-effect logistic regression, P = 0.9999). On the opposite, when prey tried to escape, probability of successful capture was 7% lower than when prey did not react (capture success = 92%, mixed-effects logistic regression, P < 0.001). The greater the flight initiation distance, the lower the probability of seals successfully capturing the prey (mixed-effects logistic regression, P < 0.001, Figure 4).

**Figure 4:**
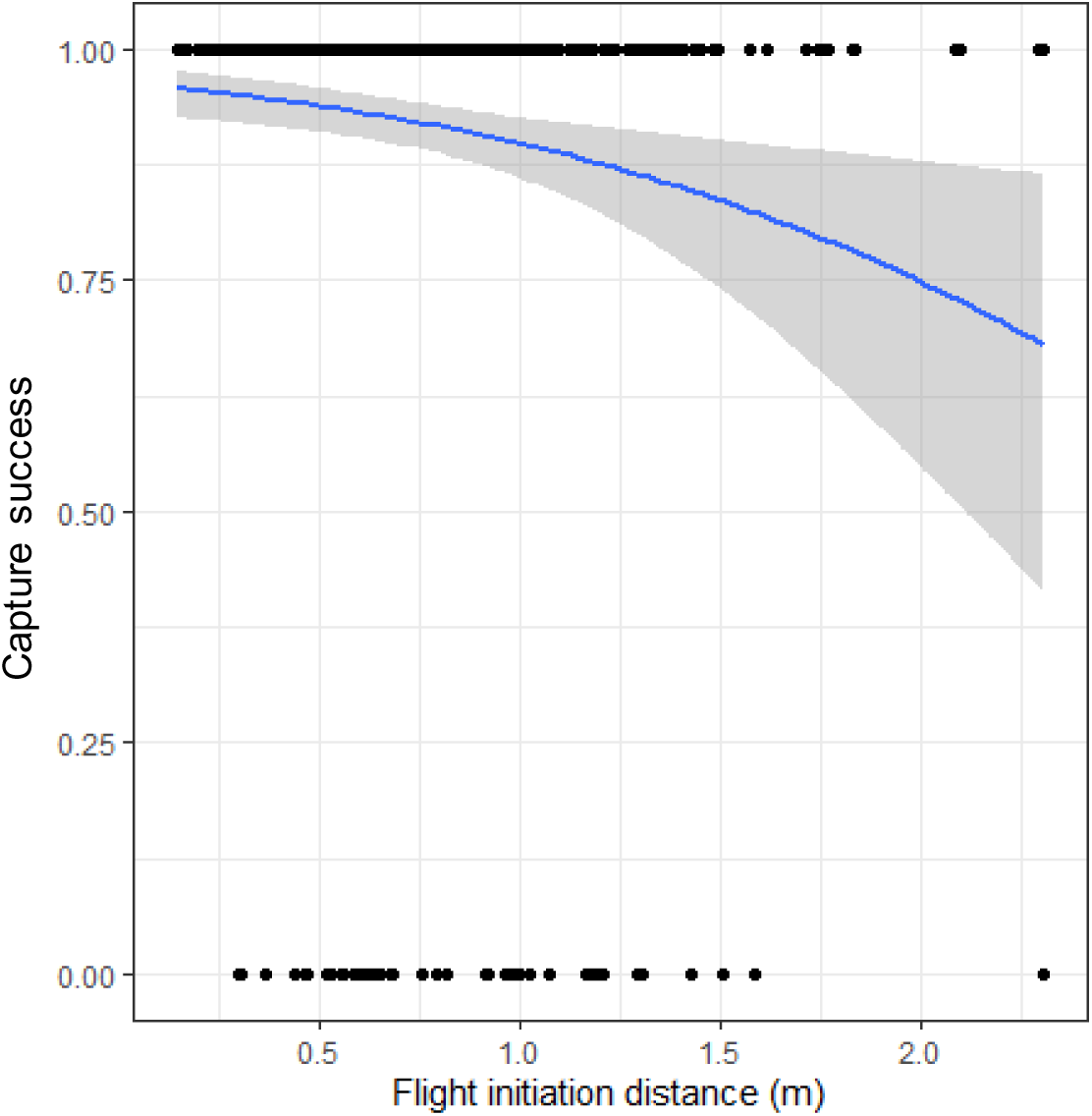
Effect of prey flight initiation distance (m) on prey capture success (%), modelled using a mixed-effects logistic regression (P < 0.001). Prey capture success was estimated from sonar recordings on 490 bouts for which prey reaction timing was inferred, for nine female Southern elephant seals equipped with high-resoluti on movement and sonar tags in Kerguelen Islands between 2018 and 2020.

## 4. DISCUSSION

We used high-resolution bioluminescence and movement sensors combined with active acoustics to study the functional relationships between Southern elephant seals and their prey. Sonar tag recordings provided novel information on characteristics (acoustic size) and fine-scale behaviour of prey targeted by seals. Tagged Southern elephant seal females from Kerguelen Islands mostly targeted small, isolated prey, while schools were almost never encountered. Prey acoustic size ranged between 4 and 10 cm, which is consistent with sizes of the main Southern elephant seals prey species (Cherel et al. 2008, Guinet et al. 2014), i.e. *Electrona antarctica* 3-10 cm, *E. carlsbergi* 4-10 cm and *Krefftichthys anderssoni* 1-8 cm (Collins et al. 2008), although it is important to remember that acoustic and physical sizes are not necessarily similar. Indeed, prey orientation has a strong influence on the measures of acoustic size (Burwen et al. 2003): if a prey is oriented perpendicular to the beam, its acoustic size reflects its width, while if it is oriented parallel to the beam, its acoustic size reflects its length. When prey tries to escape, it will at some point likely be oriented along the beam axis, so measuring the maximum length of given prey trace provides a strong indication of the true prey size. A small but meaningful proportion of prey were 10 to 20 cm (up to 35 cm) in acoustic size. these prey sizes could correspond to other abundant myctophid species in Kerguelen waters, such as *Gymnoscopelus braueri* or *G. nicholsi*, up to 13 and 19 cm in length respectively. Finally, targets larger than 20 cm could be consistent with squids, species occasionally found in elephant seals’ diet (i.e. *Martialia hyadesi* and *Todarodes filippovae* that measure up to 40 and 55 cm respectively (Cherel et al. 2008)). Further studies are needed to understand horizontal and vertical distributions of these different-sized prey in relation to environmental conditions.

High-resolution movement sensors integrated in the sonar tags also allowed us to describe the hunting sequence of free-ranging Southern elephant seals. Seals consistently started their approach by gliding towards their prey about 6 s before the capture. This is consistent with results from Chevallay et al. (in revision) that showed that elephant seals usually detect their prey between 6 and 10 s before the strike. Prey usually reacted one to two seconds before the seals’ capture, by trying to escape or by emitting a bioluminescent flash. If prey did not respond, they were captured in a few short head strikes. Larger prey likely need longer handling times and thus longer head movements (Bowen et al. 2002, Hocking et al. 2016), as already observed by Goulet et al. (submitted) in Southern elephant seals, but also in Australian fur seals *(Arctocephalus pusillus doriferus*, (Hocking et al. 2016)), sub-Antarctic fur seals (*Arctocephalus tropicalis*, (Hocking et al. 2016)) and Northern elephant seals (*Mirounga angustirostris,* (Adachi et al. 2019)) where the number of acceleration peaks correlated with prey size. If prey tried to escape, seals responded by pursuing them for several seconds, performing series of low intensity head movements before managing to capture their prey in more than 90% of the cases. Therefore, our results show that prey behaviour influences the predator’s hunting tactics. These approach phases characterised by bursts of acceleration to capture mobile prey and deceleration to capture inactive prey were also observed in other aquatic top predators such as Adelie penguins (*Pygoscelis adeliae*, (Wilson et al. 2002)) or Baikal seals (*Pusa sibirica*, (Watanabe et al. 2004)). We observed these hunting modes similarly in descents and ascents, which confirms prey-dependant behaviours in seals, rather than a consequence of the prevailing movement direction.

### Predator approach tactics: how to stay stealthy?

Seals usually stopped swimming several seconds before striking, as described by Chevallay et al. (in revision). This type of approach resembles stalking observed in terrestrial carnivorous species such as lions *(Panthera leo)*, pumas *(Puma concolor)*, or tigers *(Panthera tigris)* (Elliott et al. 1977, Sunquist 1981, Van Orsdol 1984). Stalking is defined as a slow approach without noisy body movements, allowing the predator to get very close to their prey without alerting it (Curio 2012). This kind of approach is usually observed in predators for which the costs of prey pursuit are high, as in sperm whales for example that are known to stop swimming before initiating a pursuit (Aoki et al. 2012). Elephant seals are hypometabolic predators performing long and deep dives to feed. Consequently, active hunting with long prey pursuit could result in shorter dives, as dive duration at a given depth decreases with increasing swimming effort (Génin et al. 2015). It may thus be advantageous to adopt a discreet approach that limits the risks of triggering prey escape.

Fish can detect bow waves produced by an approaching predator through their lateral lines (Blaxter & Fuiman 1990, McHenry et al. 2009, Stewart et al. 2013), a series of small mechanoreceptors located under their skins. Therefore, by adopting a slow approach, the mechanical disturbance generated by the forward movements of seals may be considerably reduced, lowering the risk of being detected by prey. Gliding before the strike was consistently observed during descending bouts. As seals are usually negatively buoyant through most, if not all the duration of their post-breeding foraging trip, they are likely to glide on descents, so they may take advantage of their inertia to approach their prey discreetly. However, gliding was also observed for 60% of strikes in ascending bouts, where seals usually swim actively. This suggests that gliding is a deliberate manoeuvre adopted by seals and not only dictated by gravity forces. The importance of slow approach to delay prey detection was highlighted in many terrestrial and aquatic species (e.g. in desert iguanas *(Dipsosaurus dorsalis)* (Cooper Jr 2003), in skinks *(Eumeces laticeps)* (Cooper Jr 1997), in Atlantic cods *(Gadus morhua)* (Meager et al. 2006), and teleost fish (Webb 1984)).

A meaningful proportion of prey (26%) showed no evidence of reaction to the seals’ approach. While the proportion of non-evasive prey may be overestimated due to the narrowness of the sonar beam (i.e. prey may have reacted outside the beam), the systematic late prey reaction supports the hypothesis that gliding behaviour may be used by seals to delay prey reaction, which may explain their very high capture success. The ontogeny of how seals acquire this specific stalking tactic is unknown. Stalking likely requires a high level of body movement control, and thus experience. Learning and experience play a key role in the ability of predators to efficiently capture their prey (Edwards & Jackson 1994, Guinet & Bouvier 1995, Morse 2000). However, unlike other carnivorous species such as felids or killer whales whose young learn by mimicking their mother or their conspecifics (e.g., (Guinet & Bouvier 1995, Elbroch et al. 2013)), elephant seals forage individually from their first day at sea. They consequently must develop their hunting tactics on their own. Equipping younger individuals with sonar and movement tags from their first year at sea could provide a better understanding of how seals develop their hunting tactics and how their hunting efficiency increases with their experience.

### Prey defence mechanisms: how not to get captured?

This study identified flash emission and escape attempt as the two types of prey reaction to predators, which could be displayed simultaneously. Escape attempt was the most frequent defence mechanism adopted by prey. Escape reaction was associated with a reduction in seals’ capture success, even though it remained high (90% of the cases). Manoeuvrability being inversely proportional to body size (Domenici 2001), small prey benefit from a greater manoeuvrability than their much larger predator. However, their swimming speed is also much lower than that of elephant seals (i.e. 0.1-0.3 m/s for myctophids (Ignatyev 1996) vs. 1-2 m/s for elephant seals (Gallon et al. 2013)). Therefore, prey must react far enough from the seals to take advantage of their higher manoeuvrability and escape capture. Indeed, we found that the greater the prey flight initiation distance, the lower the seal capture success rate.

On the opposite, flash did not seem to be an efficient defence mechanism as prey capture success was the highest for flashing prey. Flash emission is a common defence mechanism observed in marine organisms, used to temporarily dazzle predators (Barnes & Case 1974). However, myctophid flashes might be too weak to dazzle large elephant seals. Predation pressure from seal is likely lower than that from other fish and squids, for which dazzling might significantly reduce predation risk. This could explain why this flashing behaviour has been retained in fish populations even if it fails to decrease capture success. A recent study showed that elephant seals may trigger flash emission of their prey by performing a small head movement prior to capture (Goulet et al. 2020). Several studies have found that predators may benefit from flashes from their prey, e.g. Pacific angel sharks (*Squatina californica,* (Fouts & Nelson 1999)), blue sharks *(Prionace glauca,* (Tricas 1979)) and dogfish *Galeus melastomus* (Bozzanao et al. 2001)) that use bioluminescence cues to locate and capture their prey in dark areas.

This study suggests that prey characteristics and behaviours play an important role in seals’ energy expenditure and foraging efficiency. Larger prey were captured with higher swimming efforts and longer handling times, as observed in harbour seals *(Phoca vitulina*, (Bowen et al. 2002)). Similarly, approaches towards evasive prey were characterised by high swimming efforts thus likely higher energy expenditure, even if prey capture success of escaping prey remained high. In a study aiming at assessing the relationship between prey type and net energy accumulation, Goulet et al. (submitted) showed that seals foraging on evasive prey displayed a slower improvement of body condition than seals foraging on less evasive prey. Similar results were observed in double-crested cormorants *(Phalacrocorax auritus)* that have increased oxygen consumption when foraging on mobile prey compared to sedentary ones (Halsey et al. 2007).

## 5. CONCLUSION

This study offers new insight into fine-scale predator-prey interactions in Southern elephant seals. Sonar tag recordings allow for the description of the fine-scale foraging behaviour of seals and provide key information on the characteristics and behaviour of their prey. Seals’ hunting behaviour appeared to be shaped by characteristics and behaviour of their prey. Capture success was high, however, evasive prey were associated with an active pursuit that can result in significant energy costs. As prey behaviour is likely influenced by environmental conditions such as temperature or light level, elephant seals may favour foraging activity in environment where prey are less likely to react. To test this hypothesis, further studies should assess the impact of environmental conditions on prey behaviour.

## Acknowledgments

SES data were gathered as part of the “Système National d’Observation: Mammifères Echantillonneurs du Milieu Océanique” (SNO-MEMO, PI. C. Guinet) and in the framework of the ANR HYPO2. Fieldwork in Kerguelen was supported by the French Polar Institute (Institut Polaire Français Paul Emile Victor) as part of the CyclEleph programme (n. 1201, PI C. Gilbert) and with the help of the Ornitho-eco programme (n. 109, PI C. Barbraud). Data acquisition was also supported by CNES-TOSCA (Centre National d’Études Spatiales). The authors wish to thank all individuals who, over the years, have contributed to the fieldwork of deploying and recovering tags at Kerguelen Island. We also want to thank Mark Johnson for providing tags, software and codes for data analysis.

## Notes

### Competing Interest Statement

The authors have declared no competing interest.

